# Expectation effects based on newly learnt object-scene associations are modulated by spatial frequency

**DOI:** 10.1101/2025.06.02.657314

**Authors:** Morgan Kikkawa, Daniel Feuerriegel, Marta I. Garrido

## Abstract

Objects typically appear within rich visual scenes. Some models of visual system function propose that scene information is extracted from low-spatial frequency components and rapidly propagates through the visual processing hierarchy. This contextual information may help bias perceptual inferences toward objects that are likely to appear within a scene, enacted via top-down feedback carrying predictions. We tested this hypothesised influence of low spatial frequency information through newly learnt predictive object-scene associations. We recorded electroencephalographic (EEG) data from 40 participants who viewed high-spatial frequency objects embedded in either low- or high-spatial frequency scenes. Object-scene pairings were probabilistically manipulated such that certain objects more frequently appeared in certain scenes. We trained classifiers on EEG data from object-only trials and tested them on object plus scene trials. We did not observe differences in classification accuracy across expected and unexpected objects for both low- and high-spatial frequency scenes, nor any interaction between spatial frequency and expectation. However, we observed expectation effects on event-related potentials. These effects arose at similar latencies for both low- and high-spatial frequency scenes. Together, we report evidence that expectations induced by object-scene pairings influence visually evoked responses but do not modulate object representations.

## 1. Introduction

We rarely encounter visual objects in isolation. Instead, objects predominantly appear within visual scenes, which provide contextual information that aids in object recognition (Palmer 1975). For example, objects presented in ecologically valid contexts are identified faster and more accurately (Biederman 1972; Davenport and Potter 2004; Joubert et al. 2008). Although there is evidence that visual scenes can facilitate object identification, we do not fully understand how this is implemented within the brain (Joubert et al. 2008).

One proposed mechanism of scene-facilitated object recognition involves learning statistical regularities to form expectations about which objects are likely to appear in which environments (Oliva and Torralba 2007). These learnt statistical regularities are thought to influence visual processing through top-down modulations associated with predictions (Rao and Ballard 1999; Friston 2005). The predictions themselves are generated from past sensory experiences and then compared against bottom-up sensory information. Prediction errors, the discrepancy between top-down predictions and actual bottom-up input, are used to update future predictions via an iterative process implemented in hierarchically organised visual areas. A common method of instigating scene-induced expectations is to pair objects with semantically congruent and incongruent visual scenes (Biederman 1972; Davenport and Potter 2004; Joubert et al. 2008; Mudrik et al. 2010; Rossel et al. 2022; Faurite et al. 2024). For example, a car on a road is congruent, but a car in a kitchen is incongruent. Predictive processing models propose that visual scenes can assist object processing by biasing our range of perceptual inferences (Peelen et al. 2023). In this way, the subjective likelihood of an object occurring within a particular visual scene can be used to guide how incoming sensory information is interpreted. However, congruency experiments often capitalise on associations learnt over long periods of time, whereas some predictive coding models posit that statistical regularities are learnt quickly and need not be task relevant (Kok et al. 2017; de Lange et al. 2018; Richter et al. 2018; Walsh et al. 2020).

When considering scene-induced expectation effects, temporal issues may arise if we assume the visual scene must be sufficiently processed to bias perceptual inferences regarding object identity. This is because the scene and object are presented concurrently. If processing the semantic identity of the scene is required to bias the perceptual inferences relating to the embedded object, then we would expect prediction effects to manifest at later stages of visual processing, such as effects within the anterior temporal lobe (Ganis and Kutas 2003). However, given evidence of rapid visual scene processing (Potter et al. 2002; Joubert et al. 2007, 2008), models appealing to the feedforward sweep have been proposed to explain how fast scene processing can modulate object processing at earlier stages (Bullier 2001; Bar 2004). This model draws on observations that low-spatial frequency (LSF) information is processed faster than high-spatial frequency (HSF) information and leads to earlier responses within higher-order structures such as the orbitofrontal cortex (Nowak et al. 1995; Bar et al. 2006; Kveraga et al. 2007). Bar (Bar 2004) posits that LSF information from visual scenes can rapidly propagate up to higher-order areas to generate coarse sensory estimates of scene context. This contextual estimate can then influence lower-level areas, to suppress the processing of contextually improbable objects.

While it has been suggested that the feedforward sweep uses LSF estimates to guide the processing of expected objects, it is unclear whether these contextually driven expectations are the same phenomenon as described by predictive coding models (Walsh et al. 2020). Firstly, it is unclear if the feedforward sweep model applies to expectations learnt over shorter timescales. Electroencephalographic (EEG) studies have found that newly learnt object-scene and face-scene associations can produce congruency effects in event-related potential (ERP) responses (Hannula et al. 2006; Smith and Federmeier 2020, 2024). However, these studies present the scene before the object, making it difficult to assess if LSF and HSF information are used differentially to process expected and unexpected objects. Secondly, while it has been found that object identity can be decoded from fMRI BOLD responses with greater accuracy for objects in congruent scenes over objects in isolation (Brandman and Peelen 2017), it is unknown if the LSF components of the scene are responsible for influencing object representations. LSF components may induce the creation of an early sensory template for the expected stimulus, perhaps analogous to predictive pre-activation found in previous studies (Kok et al. 2017; Blom et al. 2020). Alternatively, they may produce top-down inhibition of neural activity unrelated to the processing of the expected object. This process would lead to a higher signal-to-noise ratio, resulting in a sharpened stimulus representation (de Lange et al. 2018).

Here, we assessed whether the rapid propagation of LSF information influences top-down predictions associated with newly learnt object-scene pairings. We presented objects embedded within visual scenes and entrained expectations by probabilistically pairing specific object and scene identities. Frequently paired object-scene combinations are termed expected, and infrequently paired combinations are termed unexpected. HSF objects were concurrently presented within either LSF or HSF visual scenes. We also trained support vector machines on patterns of EEG responses to HSF objects presented in isolation. We then assessed whether classification of HSF objects differed across expected and unexpected LSF and HSF scenes. We hypothesised that expectation effects on object classification would arise earlier for LSF scenes compared with HSF scenes, which would provide evidence for contextual LSF information inducing early sensory templates of expected objects. We also hypothesised a larger classification accuracy for expected objects versus unexpected objects. Finally, given the plausible bidirectional modulation of visual processing for objects and scenes (Peelen et al. 2023; Faurite et al. 2024), we also sought to assess whether spatial frequency and expectations interact for the holistic processing of object-scene pairings. This was investigated by mapping the time courses of ERP differences across conditions.

## 2. Method

### 2.1 Participants

We recruited 43 individuals who had normal or corrected-to-normal vision. Three participants were excluded due to numerous excessively noisy channels. All participants gave informed consent and received course credit for participating. Participants were aged between 18 – 29 years (*M* = 19.35, *SD* = 2.3, 31 women, 7 men, 2 non-binary). Thirty-five participants were right-handed. As interaction effects of expected object-scene pairings and spatial frequency in EEG have not been previously studied, we recruited a large sample size for EEG studies to ensure high statistical power. The study was approved by the Human Research Ethics Committee of the University of Melbourne (Ethics ID: 25177).

### 2.2 Stimuli

Stimuli were presented 80 cm from participants on a 24.5” ASUS ROG Swift PG258Q monitor (1920x1080 resolution, 60 Hz refresh rate) in a dimly lit room. Stimuli were presented using PsychoPy v2023.2.3 (Peirce et al. 2019). Code used for stimulus presentation will be available at osf.io/xgazf at the time of publication.

Visual stimuli can be seen in Fig. 1A and were greyscale images of either a dog or house (objects) presented in isolation or placed within a grey circle within a greyscale forest or canyon (scenes). Object images were taken from the THINGS database (Hebart et al. 2019). Scene images were found using the Creative Commons search engine (Whittaker 2011; Mayer 2014). Objects subtended 3° of visual angle and were high pass filtered to remove spatial frequencies below 6 cycles per degree (cpd). Objects were padded with a 0.2° grey frame buffer to mitigate edge artefacts. This padding was removed post-filtering. Scenes subtended 18° of visual angle and were either high- or low-pass filtered using cutoffs of 6 and 2 cpd, respectively. All objects and scenes were filtered using a 6^th^-order Butterworth filter (as per the Psychopy butter2d_hp and butterd2_lp functions). A grey circle of 5° visual angle in diameter was positioned in the centre of visual scenes to prevent embedded objects from overlapping with the visual scene. All objects and scenes were joint contrast normalised to reduce discriminability differences across spatial-filtering conditions (as described in Perfetto 2020). Stimuli were presented along with a fixation stimulus, which subtended 0.3° visual angle and comprised of a black circle with a negative space crosshair containing a black fixation dot at the intersection (Thaler et al. 2013).

**Figure 1.**
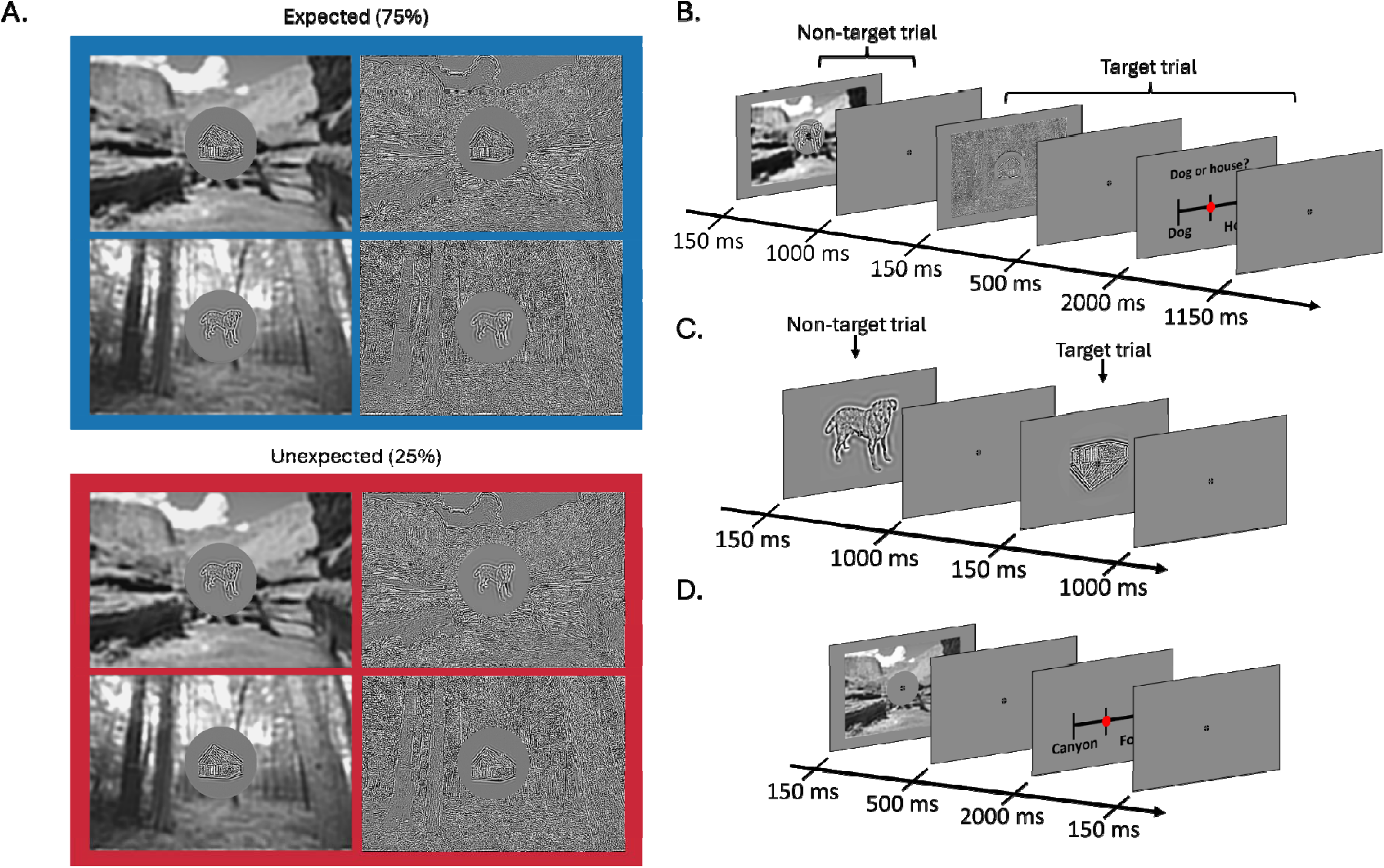
Stimuli, object-scene associations, and trial diagrams. Note that some object sizes and contrast levels have been modified for display purposes. A) Stimulus examples and object-scene associations. An HSF image of either a dog or house was placed within a 1.5° radius grey circle centred on a visual scene. Visual scenes were either a forest or canyon and could either be HSF- or LSF-filtered. Objects and scenes were paired such that a given object was more likely to occur within a certain scene. For example, across all stimuli containing the house, the house would appear within a canyon on 75% of the trials. These associations were counterbalanced across participants. B) Trial diagram for object-scene blocks. Participants viewed dogs and houses embedded within visual scenes. 10% of trials were designated target trials in which participants had to report the most recent object they had seen. C) Trial diagram for object-only trial blocks. Participants viewed an image of a dog or house in isolation and were tasked with detecting target objects (an upside-down image occurring in 10% of trials). Participants were required to give a speeded response as soon as they identified a target object. D) Trial diagram for scene-only blocks. Participants viewed both low- and high-frequency filtered scenes in isolation with a grey circle superimposed in the centre. Participants were required to identify the visual scene for every trial.

### 2.4 Experimental Design

Participants were presented with three types of blocks: object plus scene, object only, and scene only. In object plus scene trials (Fig. 1B), participants were presented with stimuli consisting of an object (dog or house) embedded within either a forest or canyon. A probabilistic pairing was made such that a dog would appear within a canyon 75% of the time, and a house would appear within a forest 75% of the time. On the remaining 25% of trials, the opposite object would appear in the visual scene. These associations were counterbalanced across participants, who were explicitly informed of the object and scene associations prior to commencing the experiment. Object and scene associations were based purely on the identity of the visual scene and not the SF content. LSF and HSF visual scenes, as well as dog and house object images, were presented equal numbers of times throughout the object plus scene blocks. Object plus scene images were presented for 150 ms following a fixation cross (same as above). The interstimulus interval was 1,000 ms for non-target trials. On 10% of trials, termed target trials, participants were tasked with identifying whether the last image they saw was a dog or house. On target trials, a response prompt screen was presented 500 ms after stimulus offset and responses were given using a TESORO Tizona numpad (1,000Hz polling rate). The prompt screen contained a dog and house label situated roughly 4° to the left and right of fixation. Participants gave a ‘1’ or ‘3’ response corresponding to the left, and right label, respectively. Each label’s position was randomised for each target trial to prevent motor preparation prior to the prompt. Participants were given 2,000 ms to respond. Response times (RT) were calculated relative to the onset of the prompt screen. Once participants provided their response, the prompt screen was displayed for another 100 ms before displaying a 1,150 ms fixation interval. These blocks contained 160 trials, totalling 1,280 trials across eight blocks. Before starting the main experimental blocks, participants first completed a practice block of 32 trials. After the practice block, participants were asked whether they remembered the scene and object associations and were reminded if they had forgotten. Of the 40 participants, only four misremembered the most likely object-scene pairings.

Blocks of object-only trials were presented in alternation with object plus scene blocks. In object-only trials (Fig. 1C), a high pass filtered image of either a dog or house was presented for 150 ms following a fixation cross. The fixation cross was presented for 1,000 ms. On 10% of trials, the presented object was upside down (target), and participants were tasked with responding to targets via a button press. Responses to target stimuli made within 700 ms were counted as hits. Trials were ordered according to a Type-1 Index-1 sequence meaning that each object preceded the other object an equal number of times. Participants performed eight blocks of this task with 153 trials per block, totalling 1,224 trials. Prior to commencing the experiment, participants also performed a practice block for the task comprising 32 trials. They were allowed to repeat this practice block as many times as they wanted until they felt comfortable with the task.

Finally, to test whether there were behavioural differences between forest and canyon scenes, participants also completed a scene discrimination task consisting of one block of 129 trials (Fig. 1D). This was conducted prior to the above blocks and involved presenting only the visual scenes with the central grey circle. Both low- and high-spatial frequency variants of the two visual scenes were used. No EEG data was collected during this task.

### 2.3 Behavioural Data Analysis

We calculated participants’ mean accuracy (proportion correct) and mean RT for each block type throughout the experimental session (excluding training). The following analyses were conducted on both mean accuracy and mean RT (using SciPy.stats). For the object plus scene blocks, we conducted one-way repeated measures ANOVAs across the expectation and spatial frequency conditions. We also performed two paired-samples t-test (two-tailed), one for expected versus unexpected, and one for LSF versus HSF. For the object-only blocks, we conducted a paired-samples t-test (two-tailed) to assess differences in identification accuracy and RT for object type (dog versus house). Lastly, for the scene-only blocks, we conducted one-way repeated measures ANOVAs to compare differences across spatial frequency and scene identity. We also performed a paired-samples t-test (two-tailed) comparing LSF and HSF scenes.

### 2.4 EEG Data Acquisition and Processing

EEG signals were recorded using a 128-channel Biosemi Active II system (Biosemi, the Netherlands) with a sampling rate of 512 Hz using common mode sense and driven right leg electrodes. Five additional electrodes were placed: two behind the ears at the mastoids, two 1 cm from the outer canthi of each eye, and 1 placed directly below the left eye.

EEG data were processed using MNE in Python 3.6 (Gramfort et al. 2013), and the dataset as well as data processing and analysis code will be available at osf.io/xgazf at the time of publication. Excessively noisy channels were first identified by determining if any channels exhibited a 500 uV peak-to-peak signal amplitude for 20% or more of either object-only or object plus scene trials. This was conducted on epoched data that was first referenced to the average of both mastoids and then high-pass filtered at 0.1 Hz and low-pass filtered at 40 Hz. The data was epoched from -100 to 500 ms relative to stimulus onset. Noisy channels were then excluded from the average reference calculations and independent components analysis (ICA). The original dataset (with bad channels removed) was then referenced to the average of all channels and filtered (using 0.1 and 40 Hz cutoffs). Electrode AFz was removed to account for the rank deficiency caused by average referencing and ICA was performed using the infomax algorithm (Lee et al. 1999). Independent components were then manually inspected to identify blink and saccade artefacts following guidelines by Chaumon and colleagues (Chaumon et al. 2015). These components were removed from the filtered dataset and a final sweep of bad channels was conducted with a more sensitive criterion. Bad channels were identified in the same manner as before but with a 100 µV peak-to-peak cutoff. Identified noisy channels and AFz were then interpolated using the spherical spline method (Perrin et al., 1988). The processed data was then epoched from -100 to 500 ms relative to stimulus onset and baseline corrected from -100 to 0 ms. Trials were rejected if any one channel displayed a peak-to-peak amplitude greater than 150 µV during the epoched window.

### 2.5 Multivariate Pattern Analyses (MVPA)

We used linear support vector machines (as implemented in SciKitLearn, version 1.6.1, kernel = linear, and default) to assess whether object classification performance differed across expected and unexpected object-scene pairings. First, to verify that object identity could be decoded, models were trained and tested using data from object-only blocks. A sliding window average of 28 ms in 4 ms steps was applied to the data. A 20-fold leave-one-out cross-validation method was used such that the dataset was divided randomly into 20 folds, trained on 19 and then tested on the remaining fold. This was repeated 20 times until every trial had been used for training and testing. Classification accuracy (measured as percentage of trials correctly identified by the model) for each time point was calculated as the average across folds. This procedure was repeated using every combination of time points for training and testing to produce a temporal generalisation matrix (King and Dehaene 2014).

Above-chance decoding for the object-only trials was calculated by generating a permuted labels classification performance and comparing it with the observed classification performance. The permuted labels performance represents the level of decoding that can be expected from chance. The permuted labels condition was generated by testing and training support vector machine models on neural data in which the trial labels were randomly shuffled. This was repeated 10 times for each participant to ensure no single set of random permuted label assignments biased chance level classification. The permuted labels classification performance for each participant was taken as the average of their 10 shuffles. Above-chance classification was then assessed using a cluster-based permutation test which compared observed classification and permuted labels classification performance (Maris and Oostenveld 2007; for details see below).

Once it was verified that the object identity could be classified at above-chance levels, we then tested if expectation and spatial frequency influenced the classification of object identity. We trained support vector machine models on object-only trials and tested them separately on the four types of object plus scene trials (LSF expected, LSF unexpected, HSF expected, and HSF unexpected). Both the training and testing data was averaged using the same sliding window procedure as before and the model was trained and tested on every combination of time points to once again produce temporal generalisation matrices. An effect of expectation was assessed within each spatial frequency condition by using a cluster-based permutation test between conditions (LSF expected versus LSF unexpected, and HSF expected versus HSF unexpected). We then used a cluster-based permutation test to assess the presence of an interaction effect. This was done by comparing the difference in classification differences of expected and unexpected trials between LSF and HSF. As interaction effects can occur even in the absence of main effects, we decided a priori to conduct this test regardless of whether we observed any main effects.

### 2.6 ERP Analyses

ERPs for each condition were calculated by averaging all trials within each condition for each participant. Like our MVPA, this resulted in four ERP conditions: LSF expected, LSF unexpected, HSF expected, and HSF unexpected. As expectation conditions would include neural responses to trials containing dog and house stimuli, we ensured that each object had an equal contribution to the overall ERP of the condition. To do this, the ERP evoked by each object was first calculated separately for every condition, and then the two ERPs (one for the dog, and one for the house) were averaged together. A mass-univariate analysis was then conducted between ERP conditions at a within-participants level using a cluster-based permutation test. We made three ERP comparisons: LSF expected and LSF unexpected, HSF expected and HSF unexpected, and LSF unexpected minus LSF expected, and HSF unexpected minus HSF expected. These tests allowed us to identify spatiotemporal clusters displaying significant differences.

### 2.7 Cluster-based Permutation Testing

To assess significant differences in both MVPA and ERPs, we used cluster-based permutation tests at a within-participant level. For every comparison, a null distribution was made using 5,000 permuted samples. For each sample, the condition labels of a random subset of participants were swapped, and a paired-samples t-test was performed for every time point, producing a *t*-value. *t*-values corresponding to an uncorrected *p*-value of 0.01 or less were taken as cluster-forming time points, and all adjacent time points exhibiting *t*-values over the threshold were grouped together to form clusters. For the ERP data, adjacency was defined spatiotemporally. The sum of all *t*-values within a given cluster was used to calculate its cluster mass. For each of the 5,000 permutation samples, the largest cluster mass was taken to estimate the null distribution. Finally, to identify statistically significant clusters within the observed data, cluster masses for the non-permuted condition labels were calculated and compared with the null distribution of cluster mass statistics. This allowed for the p-value of every observed cluster mass to be calculated using its percentile ranking within the null distribution. This comparison was two-tailed with any clusters exhibiting *t*-values over a family-wise alpha of 0.05 being significant.

## 3. Results

### 3.1 Behavioural Results

We first assessed if specific scene identities displayed different discriminabilities by performing repeated measures ANOVAs on accuracy scores and mean RTs from scene-only trials (Fig. 2A, 2B). Neither accuracy (*F*(3, 36) = 1.55, *p* = .20), nor RT (*F*(3, 36) = 0.44, *p* = .73) were significantly different across any of the conditions. We also assessed whether the spatial filtering of the scene affected its discriminability by conducting a paired samples t-test (two-tailed) comparing low- and high-spatially filtered scenes (Fig. 2C, 2D). Accuracy scores were found to be significantly higher for the low-spatial frequency condition (t(39) = -2.79, p =.008). However, no significant differences were found for RTs (t(39) = 1.87, p = .067).

**Figure 2.**
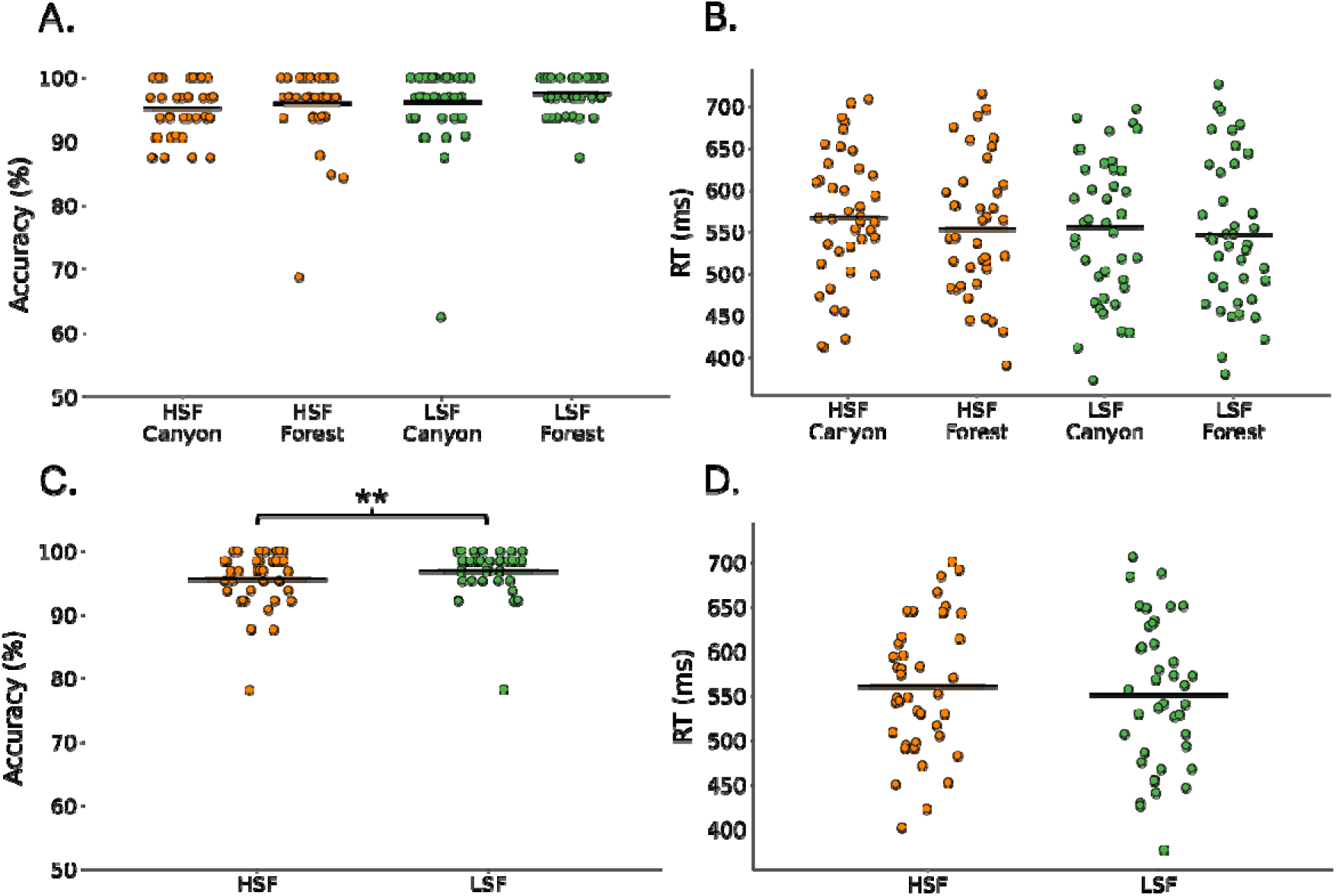
Behavioural task performance for scene-only identification task. Black horizontal lines represent the group mean. ** denotes *p* < .01. A) Accuracy by scene identity and spatial frequency type. B) Mean RTs by scene and spatial frequency type. RTs were calculated relative to the onset of the response screen which appeared 500 ms after the offset of the image. C) Accuracy by spatial frequency type. D) RT by spatial frequency type.

The mean accuracy and mean RTs for the object only identification task was also compared using a paired samples t-test (two-tailed; Fig. S1). RTs for house targets were significantly slower compared with dog targets (t(39) = 3.57, p < .001) but there were no significant differences in accuracy (t(39) = 1.38, p = .18).

Finally, we assessed differences across target trials for the object plus scene condition (Fig. 3). Trials were grouped together based on expectation and SF conditions, yielding four conditions: LSF expected, HSF expected, LSF unexpected, HSF unexpected (Fig. 3A, B). One-way repeated measures ANOVAs were performed and did not reveal significant differences in accuracy (*F*(36) = 0.44, *p* = .72) nor reaction time (*F*(36) = 0.18, *p* = .91). When grouped by expectation (Fig. 3C, D), no significant difference in accuracy was found between the two conditions (*t*(39) = 0.89, *p* = .38) but the unexpected condition displayed significantly faster RTs (*t*(39) = 2.10, *p* = .04). No significant differences were found between spatial frequency conditions for accuracy (*t*(39) = -0.55, *p* = .59) or RTs (*t*(39) = 0.99, *p* = .33).

**Figure 3.**
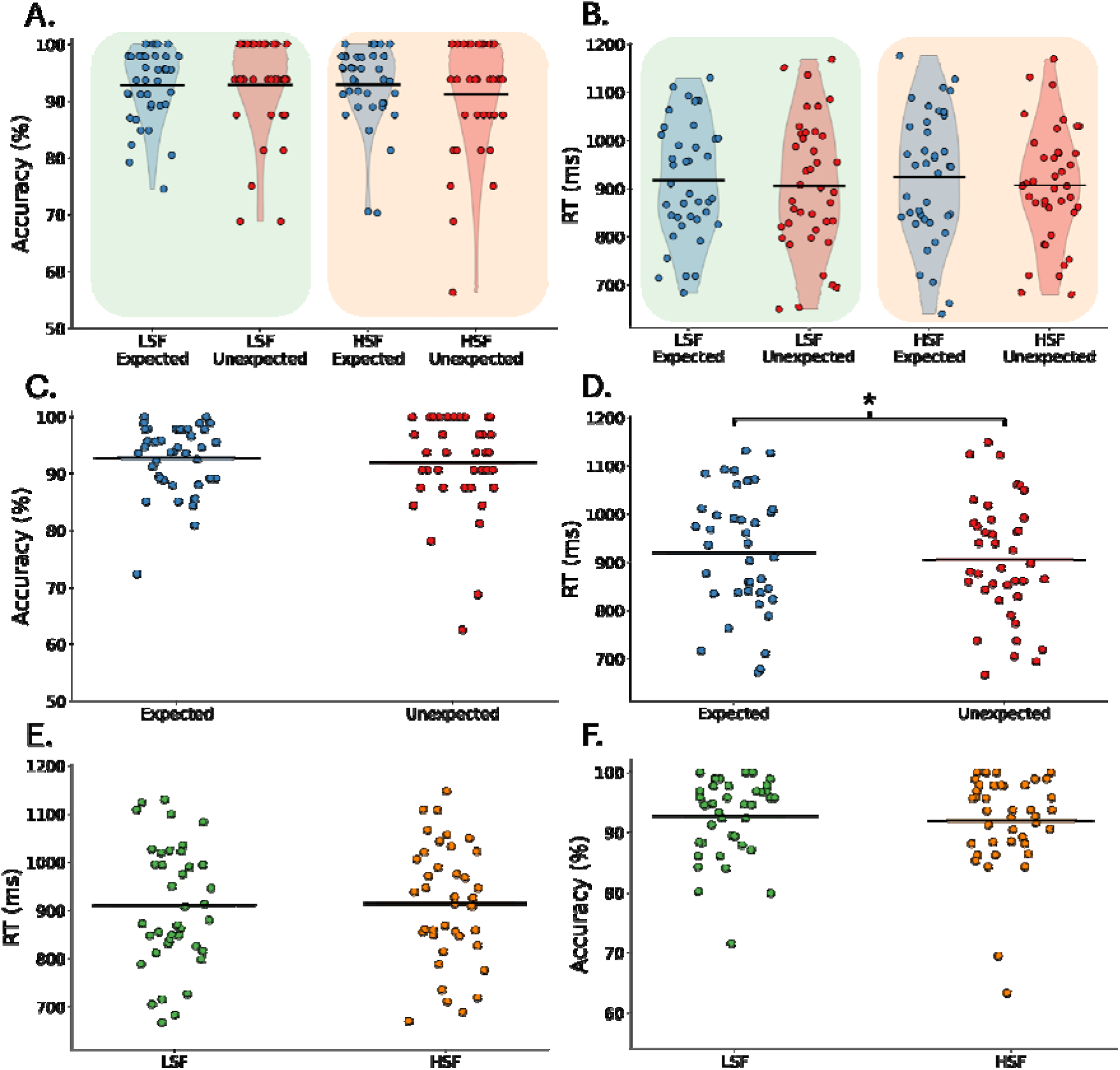
Behavioural task performance for the object plus scene task. Black horizontal lines represent the group mean. * denotes significance of p < .05. A) Mean RTs by expectation and spatial frequency condition. RTs were calculated relative to the response screen onset which appeared 500 ms after the offset of the stimulus. B) Accuracy for each participant by expectation and spatial frequency condition. C) Mean RTs by object-scene expectation. D) Accuracy for each participant by object-scene expectation. E) Mean RTs for each participant by scene spatial frequency. F) Mean accuracy for each participant by scene spatial frequency.

### 3.2 Object Image Classification

We assessed whether classification of object images based on EEG signals differed across expected and unexpected object-scene pairings, and whether the spatial frequency content of the scene modulated these effects. We first determined that the object images (when presented in isolation) could be classified at above-chance levels. We observed above-chance classification performance from ∼62 ms after image onset, which peaked at around 129 ms and lasted until 500 ms (i.e., the end of the analysed time range, results displayed in Fig. S2.).

Next, we determined if classifiers trained using EEG responses to objects in isolation could successfully classify objects that were embedded within scenes. We found significant above-chance decoding with an onset ranging from 74 – 86 ms across conditions (Fig 4A.). This extended to 500 ms for all but the HSF unexpected condition (i.e., the end of the analysed time range). We did not identify any clusters of statistically significant differences when comparing expected and unexpected conditions for either low- or high-spatial frequency scenes (Fig 4B). As it is possible for an interaction effect to occur even when both measured differences are not significant (known as a cross-over interaction), we performed another cluster-based permutation test to test for the presence of an interaction effect. This test compared the classification accuracy differences across expected and unexpected conditions within LSF and HSF conditions (Fig 4B). This analysis likewise revealed no statistically significant interaction effects.

**Figure 4.**
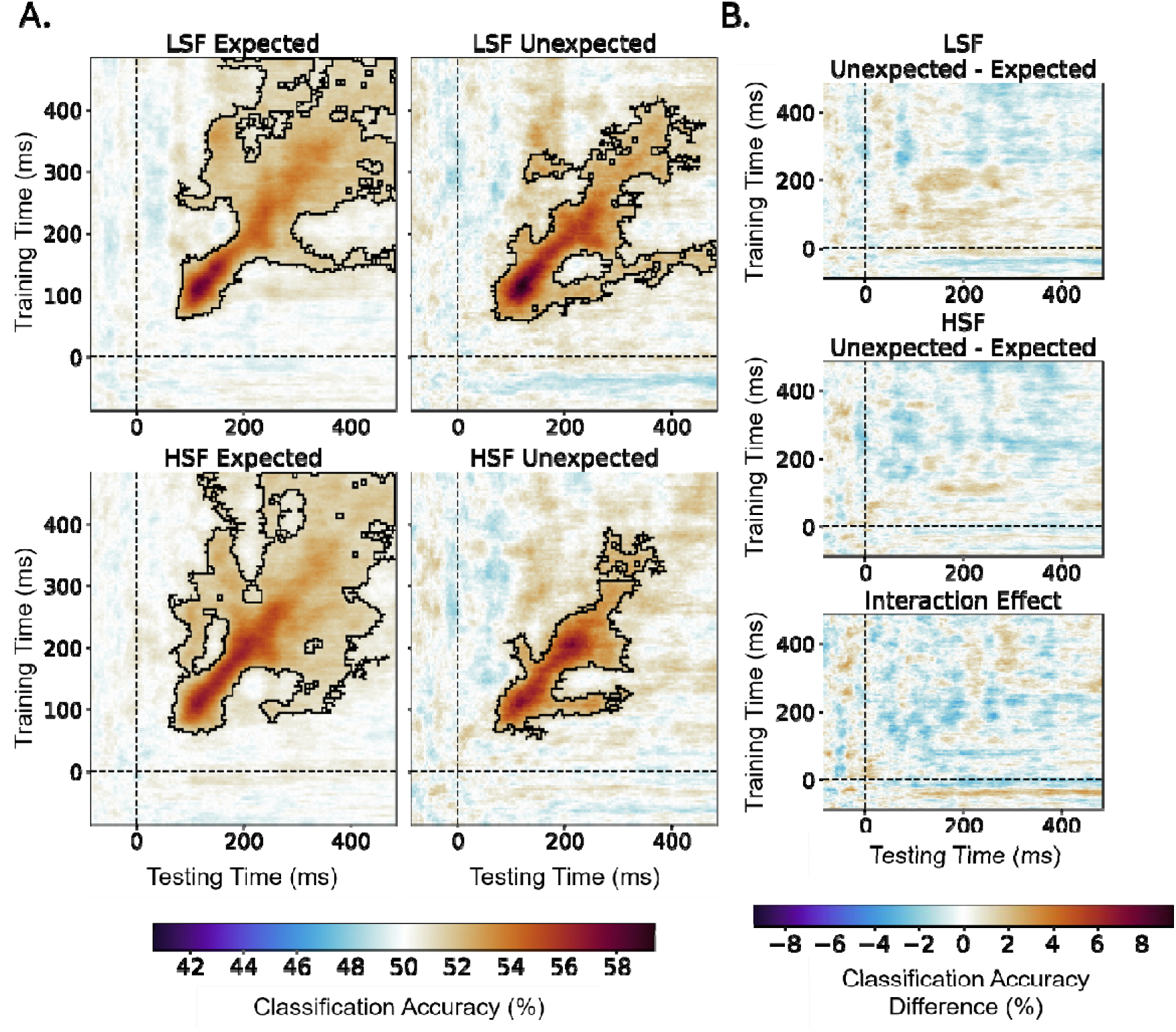
Temporal generalisation matrices of embedded object classification performance. Training times points correspond to neural responses elicited by object-only trials. Testing times correspond to the neural responses elicited by object plus scene trials. Dashed black lines represent stimulus onset. Significant clusters were determined using permutation testing and are denoted by black borders (cluster p < .05). A) Object classification performance by expectation and spatial frequency condition. Significance was determined by comparing observed classification with a permuted labels classification performance. B) Classification differences by spatial frequency condition and interaction effect. Classification accuracy differences were calculated by subtracting the classification accuracy of the expected condition from the unexpected condition. Significance was determined through a cluster-based permutation test comparing the unexpected and expected conditions within each spatial frequency separately. We then tested for an interaction effect by comparing the expectation differences of LSF and HSF conditions. The TGM was generated from subtracting the top plot (LSF Unexpected – Expected) from the middle plot (HSF Unexpected – Expected).

### 3.3 ERPs Across Expectation and Spatial Frequency Conditions

We next assessed effects of expectation on ERPs. While the classification analyses showed no significant differences, the support vector machines were trained on object-only trials and were thus designed to detect differences in neural activity relating only to the object, and not the scene. As the ERPs reflect the responses to the entire stimulus, that is, the object and the scene, differences may occur in the overall responses. We performed mass-univariate analyses and found expectation effects for both low- and high-spatial frequency conditions, as well as an interaction effect between spatial frequency and expectation.

#### 3.3.1 LSF Expectation effects

We found early effects of expectation on ERP responses to object-scene stimuli for the LSF condition, arising from 68 ms relative to stimulus onset. These effects lasted until the end of the epoched window (500 ms), occurring primarily at posterior electrodes distributed across central, centro-parietal, and parietal regions (Fig. 5A). These effects were found across three significant clusters. To better topologically visualise the effects, four time windows showing stable effects were manually chosen. Window averaged t-values are shown in Fig. 5B. Earlier expectation effects occurred between 68 – 142 ms and were focused over centro-parietal and occipital electrodes. This then consolidated to the midline and some bilateral temporal regions from 187 – 257 ms, then to fronto-central electrodes (261 – 376 ms), and finally to central and temporal electrodes (380 – 500 ms).

**Figure. 5.**
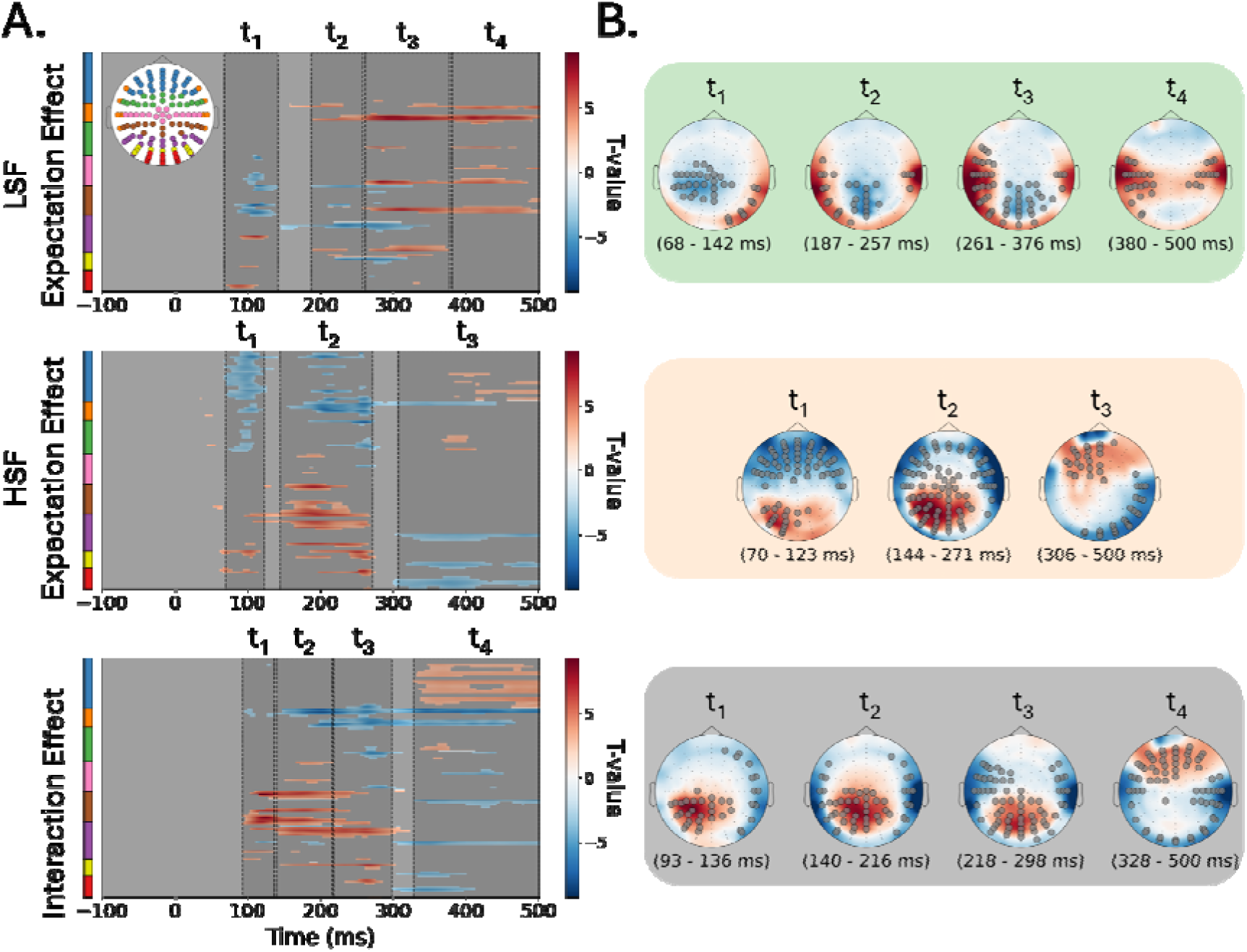
Expectation effects for low- and high-spatial frequency scene conditions. A) Timepoint-by-electrode matrix of statistically significant ERP differences. A colour bar has been provided for each matrix to designate channel location, corresponding to the scalp map icon at the top left corner. Red signifies more positive-going ERP amplitudes for the unexpected compared to expected object-scene conditions. For the interaction effect, red corresponds to the HSF expectation difference being more positive-going than the LSF expectation difference. Grey corresponds to non-significant effects. Shaded regions between dashed black lines correspond to time windows used to generate scalp maps in (B). B) Scalp maps of expectation effects by spatial frequency condition. Each scalp map is the average t-value occurring in the accompanying time window shown in (A). Values are unthresholded. Statistically significant electrodes are marked with grey circles and electrode positions are marked with small black dots.

To better visualise how the underlying activity of electrodes were modulated by expectation, we created averaged ERPs for each cluster of electrodes (Fig. 6). We found that the direction of expectation effects changed depending on the cluster, with two unilateral clusters in temporal regions exhibiting more positive-going ERPs for the expected condition. Conversely, one cluster, centred on central and centro-parietal areas showed more positive-going waveforms for the unexpected condition.

**Figure 6.**
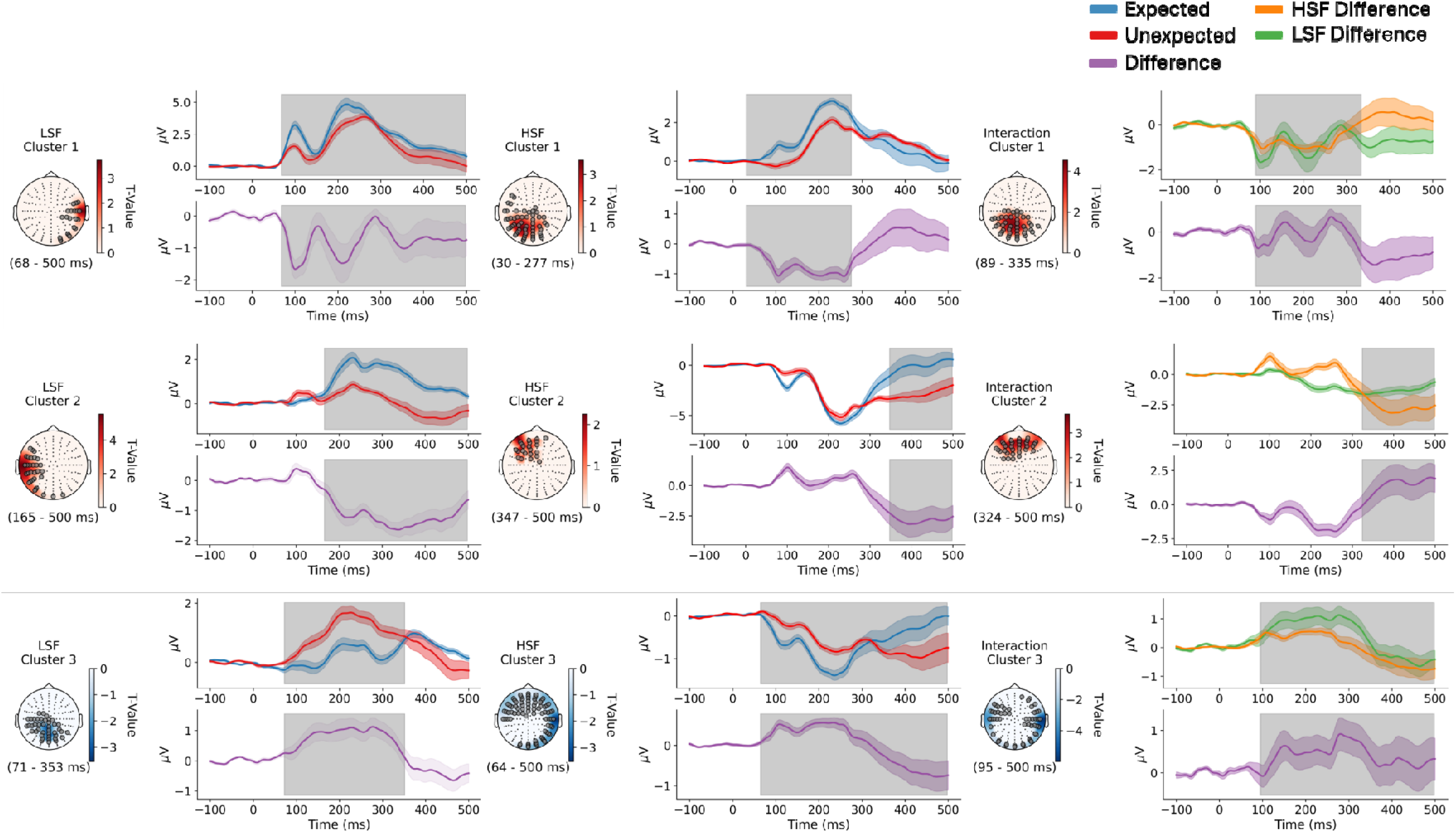
Scalp maps and group average ERPs displaying expectation and interaction effects. Each cluster has been plotted as a scalp map showing the electrodes exhibiting statistically significant effects (left) and ERPs of each condition along with a difference plot between the two conditions (right). Scalp maps display t-values averaged across the cluster time window and are thresholded such that only significant electrodes are included, indicated by grey dots. ERPs were calculated as the average activity across all electrodes in each accompanying cluster. Standard errors of the mean are displayed as coloured shading. Grey shading indicates significant differences between conditions. Interaction effect ERPs were calculated as the difference between unexpected-expected condition differences for low compared to high spatial frequency scenes.

#### 3.3.2 HSF Expectation effects

An early effect of expectation on ERP responses was also found for the HSF scene condition. We identified three significant clusters with expectation effects ranging from 30 – 500 ms relative to stimulus onset (Fig. 5A). These effects were broadly categorised into three time windows for visualisation (Fig. 5B, middle). Compared to the LSF condition, early differences occurred unilaterally over more posterior electrodes (70 – 123 ms), with a large distribution of frontal electrodes also displaying significant differences. Unlike the LSF condition, posterior regions displayed significantly more positive-going potentials for the expected condition. These effects became even more concentrated in more central electrodes at around 144 – 271 ms and then were apparent at frontal electrodes at later time stages (306 – 500 ms).

When assessing ERPs averaged over electrodes within significant clusters, like the LSF condition, the direction of expectation effects varied across clusters (Fig. 6). For the first HSF cluster, the expected condition yielded more positive-going ERPs than the unexpected condition. This cluster had a similar topography and timing as cluster three of the LSF condition. Therefore, a set of electrodes exhibited differential expectation effects dependent on the spatial frequency of the scene. A second cluster of frontal electrodes was identified, which exhibited a more negative-going potential for the unexpected condition, occurring from 347 ms.

## 4. Discussion

We assessed whether LSF information present in visual scenes facilitates top-down predictions about the identities of object embedded within those scenes. To do this, we tested for expectation effects induced by newly learnt object-scene associations, as well as their modulation by the spatial frequency content of the scene images. Contrary to our hypotheses, we found no evidence for expectation influencing the classification of objects embedded in either LSF or HSF scenes. However, we observed differences in ERPs evoked by expected compared to unexpected objects. Notably, these ERP expectation effects differed across LSF and HSF visual scenes. Our findings indicate that the information provided by visual scenes for newly learnt object-scene pairings does not modulate the early (<200 ms) visual processing of objects. However, they do produce expectation effects either through holistically modulating the response to the scene plus object stimulus, or via an object-induced modulation of responses to the visual scene. However, we did not find evidence for the type of LSF-facilitated predictions that are described in feedforward sweep models of visual processing (Bullier 2001; Bar 2004).

### 4.1 Expectations Based on Object-scene Pairings do not Influence Object Decodability

No studies to date have addressed whether newly learnt statistical regularities between concurrently presented objects and scenes can lead to differences in object classification accuracy. This makes direct comparisons with previous studies difficult due to varying methodological differences. However, our results are broadly inconsistent with previous literature. Experiments by Brandman and Peelen (2017) showed that objects presented within degraded contextual scenes are classified with higher accuracy compared to objects presented in isolation. Their study only included a congruent condition, meaning no comparison was made between expected and unexpected objects. Other MVPA studies have shown that objects appearing in their canonical locations within the visual field also elicit higher decoding accuracy, or that object-scene pairings can be decoded based on their congruency label (Draschkow et al. 2018; Kaiser and Cichy 2018).

One explanation for our lack of classification differences between expectation conditions could be attributed to the time required to learn statistical regularities. For example, congruent pairings often use objects and scenes with strong semantic and/or spatial relationships assumed to be learnt over participants’ lifetimes. These reflect deeply learnt priors that could be embedded within the visual system and produce stronger differential neural responses (Teufel and Fletcher 2020). In contrast to congruency, our study relies on object-scene pairings learnt within an experimental session that are assumed to have a neutral semantic association. This allowed us to investigate expectations as described in predictive coding models, which posit the emergence of predictions over both long and short timescales (de Lange et al. 2018; Walsh et al. 2020). While these models do not arbitrate the types of predictions learnt over short periods, the present results suggest that scene-induced expectation effects on object processing may arise over longer periods of repeated exposure, rather than within experimental time frames as done here.

Some studies have shown that statistical regularities learnt within single experimental sessions can produce differences in the decoding of visual stimuli (Kok et al. 2012, 2017; Moore et al. 2024), but findings are mixed (Richter et al. 2018; den Ouden et al. 2024). These experiments employed probabilistic cueing designs that involved the sequential presentation of a cue followed by a stimulus, whose most likely identity is predicted by the cue. This sequential presentation may be an important factor when addressing the learning time scales required for observing expectation effects. Past studies suggest that newly learnt contextual associations require cue-target delays (Smith and Federmeier 2024). Alternatively, onset asynchronies may be important for all scene-induced expectation effects, newly learnt or otherwise. In line with past work, our findings support the notion that the visual system may require time to accumulate evidence from scene contexts to produce predictions aiding in object processing (Roux-Sibilon et al. 2019).

### 4.2 Expectations Based on Object-scene Pairings Modulate ERPs

Despite no observed differences in object classification performance, we report ERP differences across expectation conditions for both LSF and HSF scenes. We also observed an interaction effect between expectation and SF. The latencies of expectation effects by SF condition were comparable, with HSF expectation effects arising slightly earlier than LSF effects. Given the feed-forward sweep posits LSF information propagates faster than HSF information, we expected that LSF scenes should induce earlier differences across expectation conditions (Bar 2004). These findings suggest that early sensory processing may not be modulated by contextual information solely from LSF domain. However, due to the inherent limitations of inferring exact latencies from cluster-based approaches, and the small latency difference observed, it is difficult to comment on the precise time courses of these effects (Maris and Oostenveld 2007).

The ERP differences we observed may reflect how objects influence the processing of visual scenes. We can arrive at this conclusion by considering the discrepant ERP and MVPA findings. Our support vector machines were trained on object-only responses and were designed to detect differences in object-selective neural responses. The mass-univariate ERP analysis detected changes in neural responses elicited by both the scene and embedded object. If expectation effects are seen only at the ERP level, the modulation may be related to the processing of the scene, and not the object. Indeed, past studies have identified effects of embedded objects influencing the processing of scenes (Lukavský 2019; Brandman and Peelen 2023; Faurite et al. 2024). These effects could be particularly relevant when scene information is less discriminable, as was the case for the HSF scene stimuli. This is further supported by lower accuracy for HSF over LSF scenes in the scene discrimination task (Fig. 2D). Therefore, it may be the case that the direction of object-scene facilitation may be governed by the relative visual evidence available from each component. This is in line with models proposed by Peelen and colleagues (Peelen et al. 2023) which conceptualise object and scene processing as a form of non-hierarchical Bayesian inference. Crucially, they propose that both the direction and magnitude of object-scene interactions are governed by weighting the relative reliabilities of the visual information. In our experiment, objects may have provided the most reliable source of sensory information, and hence, assisted in the processing of the scene. Future studies could assess object-induced scene facilitation by training machine learning classifiers on scene-only trials to determine whether expected objects boost the classification performance of scene identities. Furthermore, we would expect this increase in classification to be greater for HSF scenes compared to LSF scenes.

The early latencies of expectation effects for both LSF and HSF conditions suggest that expectation effects in object-scene pairings modulate early visual processing (∼70 ms). This is contrary to findings from Ganis and Kutas (2003) who report no ERP congruency effects before 300 ms. A number of studies have also found late ERP effects of congruity but these studies use pre-defined windows (Mudrik et al. 2010; Demiral et al. 2012; Dyck and Brodeur 2015; Smith and Federmeier 2020, 2024). However, Ganis and Kutas (2003) presented visual scenes 300 ms prior to the object, allowing more time for the visual scene to be processed. Given this, the reported ERP differences in the latter papers may not reflect object-induced influences on scene processing, but higher-level properties or associations evoked by the scenes. As we present the scene and object concurrently, the influence of objects on scene processing may not be limited to later ERP components.

### 4.3 Limitations

Our ERPs differed across a dispersed range of electrodes and time points. While we could infer the presence of expectation effects, we did not determine the cortical origins of these effects. In the absence of electrode digitisation and individual structural MRI scans, our data is not ideal for applying source reconstruction. That said, our central aim was to assess whether LSF scenes induce earlier expectation differences for embedded objects compared with HSF scenes. While our MVPA methods were well-suited to address this question, our ERP analysis was limited when considering the bidirectional influence of objects and visual scenes.

Another limitation of our study was the design of the stimuli. HSF filtering often leads to reductions in contrast, making them less distinguishable than their LSF counterparts (Perfetto et al. 2020). To address this, we performed joint contrast normalisation on LSF and HSF images to equalise discriminability. A caveat of joint contrast normalisation, however, is that it introduces residual LSF information (Kauffmann, Chauvin, et al. 2015; Kauffmann, Ramanoël, et al. 2015). Hence, our HSF stimuli may have contained some LSF information that in turn may have been sufficient to activate early sensory templates of embedded objects. This could potentially explain the lack of temporal differences found between the two SF conditions. However, the LSF images still contained substantially more LSF information than the HSF images and therefore should provide more contextual information capable of generating predictions. Arguments could be made as to the minimum amount of LSF information or the SF cutoffs required for earlier activation; however, this is an ongoing debate and not a key question of the study (for a review on the separability of LSF and HSF pathways, see Edwards et al., 2021).

Lastly, we did not behaviourally validate that participants were using object-scene associations for perceptual decision-making. This is because we did not use a speeded response and instead included a 500 ms response delay for target trials. This limits the conclusions that can be drawn in relation to specific predictive coding accounts that suggest neuroimaging effects of expectation are only observable when participants must rely on contextual information to complete the task (Richter and De Lange 2019; Alink and Blank 2021). However, our hypotheses address more general theories of prediction that assert expectation effects arise automatically, regardless of behaviour, as found by previous studies (den Ouden et al. 2009; Kok et al. 2012, 2017). Additionally, all participants were directly informed of the pairings, and, like previous studies, we assume participants learn the presented associations through large amounts of repeated exposure to object and scene combinations (Tang et al. 2018; Moore et al. 2024).

### 4.4 Conclusion

Our findings indicate that object-selective visual representations may not be influenced by information present in visual scenes or statistical regularities learnt over shorter periods of time. More specifically, we did not find evidence of contextual LSF information activating sensory templates of expected objects earlier than HSF information. Instead, it is likely our paradigm led to expected objects modulating the early sensory processing of concurrently presented scenes, reflected as differences in distributed patterns of ERPs. We conclude from this that the direction of modulation between newly learnt object-scene pairings may be governed by their relative discriminability. Sensory components containing the most available visual evidence may facilitate the processing of components with less visual evidence. We propose our results support models of object-scene processing that posit a bidirectional influence of object-scene interactions governed by sensory reliability.

## Acknowledgements

This work was supported by the Melbourne School of Psychological Sciences Studentship and the Australian Government Research Training Program. Funding sources had no role in study design, data collection, analysis or interpretation of results. The research was also supported by The University of Melbourne’s Research Computing Services and the Petascale Campus Initiative.

