## Supplementary Figures for "Expectation effects based on newly learnt object-scene associations are modulated by spatial frequency"

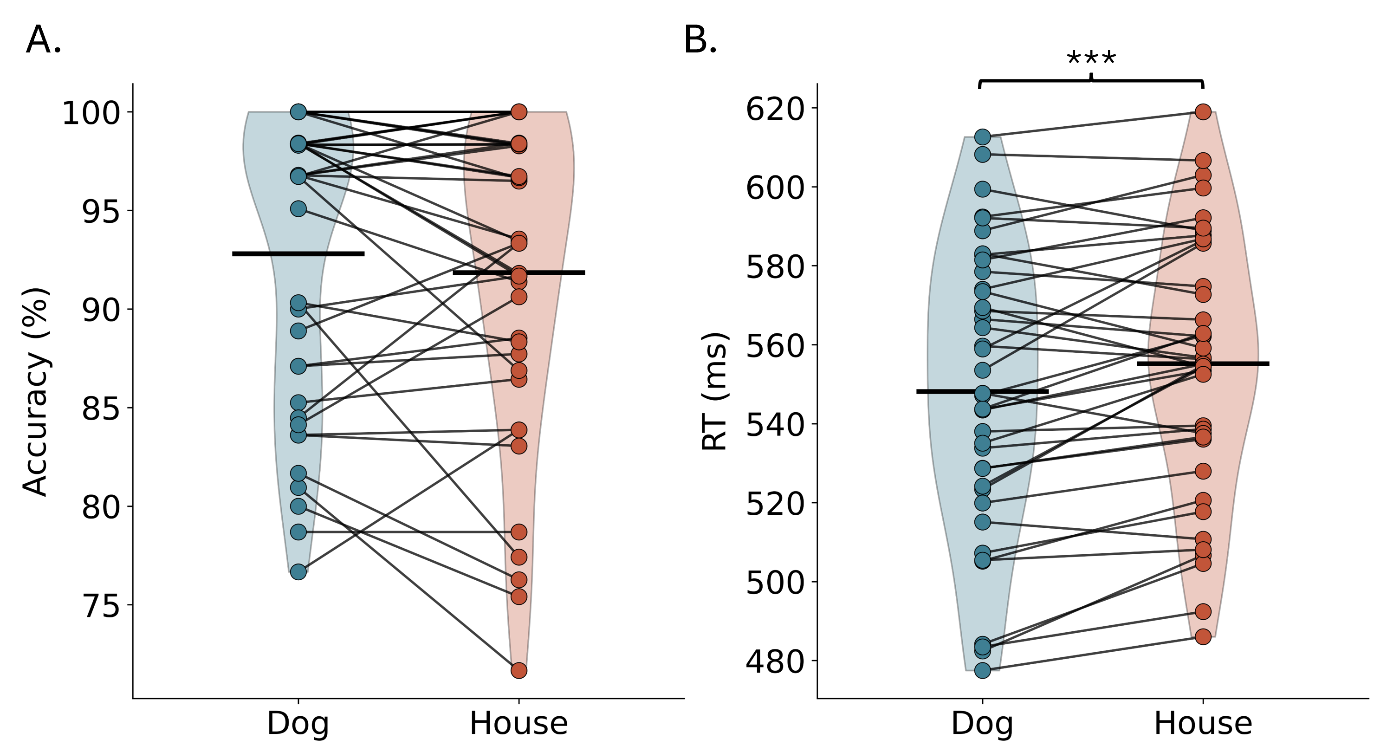

**Figure S1.** Performance in the object only identification task. A) Accuracy by object identity. B) Mean response times (RTs) by object identity. RTs were calculated relative to the onset of each upside down (target) image. Black horizontal lines represent group means. *** denotes *p* < .001.

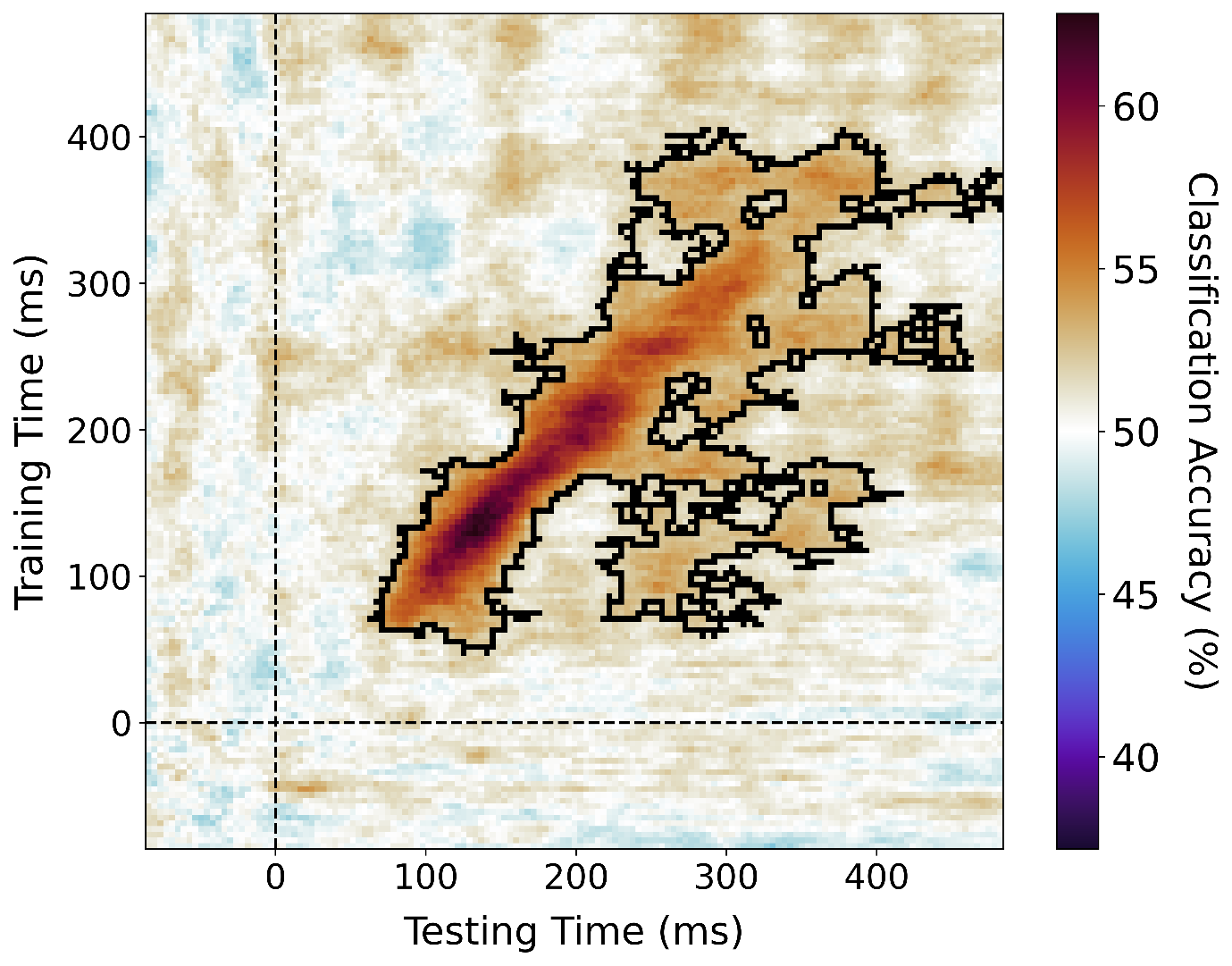

**Figure S2.** Temporal generalisation matrix showing classification performance in object-only trials. Classification accuracy was averaged across all participants. Training and testing times correspond to neural responses elicited by object only trials. A 20-fold leave-one-out cross-validation approach was used. Classification performance was calculated as the average accuracy score across all folds. Dashed black lines represent stimulus onset. Statistically significant clusters were determined using cluster-based permutation tests and are denoted by black borders (cluster *p* < .05).
